# Allosteric Modulation of Fluorescence Revealed by Hydrogen Bond Dynamics in a Genetically Encoded Maltose Biosensor

**DOI:** 10.1101/2023.12.20.572600

**Authors:** Melike Berksoz, Canan Atilgan

## Abstract

Genetically encoded fluorescent biosensors (GEFBs) proved to be reliable tracers for many metabolites and cellular processes. In the simplest case, a fluorescent protein (FP) is genetically fused to a sensing protein which undergoes a conformational change upon ligand binding. This drives a rearrangement in the chromophore environment and changes the spectral properties of the FP. Structural determinants of successful biosensors are revealed only in hindsight when the crystal structures of both ligand-bound and ligand-free forms are available. This makes the development of new biosensors for desired analytes a long trial-and-error process. In the current study, we conducted µs-long all atom molecular dynamics (MD) simulations of a maltose biosensor in both the *apo* (dark) and *holo* (bright) forms. We performed detailed hydrogen bond occupancy analyses to shed light on the mechanism of ligand induced conformational change in the sensor protein and its allosteric effect on the chromophore environment. We find that two strong indicators for distinguishing bright and dark states of biosensors are due to substantial changes in hydrogen bond dynamics in the system and solvent accessibility of the chromophore.

## 1. Introduction

Genetically encoded fluorescent biosensors (GEFBs) are protein-based indicators that can be used for dynamic imaging of practically any molecule or biochemical process in living cells. Broadly, GEFBs have a modular structure whereby a fluorescent reporter is genetically fused to a sensing domain. They fall under two main categories; Förster resonance energy transfer (FRET)-based biosensors which consist of two fluorophores (donor and acceptor) linked by a sensing domain and single fluorescent protein (FP) based biosensors where a fluorescent protein is linked to a ligand binding protein. FRET based biosensors usually require a sensing domain that undergoes a significant conformational change upon ligand binding that effectively changes the distance between the two fluorescent proteins.^1^ In single FP sensors, communication between the two modules is usually achieved by inserting a circularly permuted FP into the sensing domain. Circular permutation of FPs provides a unique opportunity to bring the chromophore, which is responsible for the spectral properties, to close the proximity of the sensing domain.^2^

In a seminal paper, it has been shown that yellow FP preserves its fluorescence when Y145 is replaced with calmodulin (CaM) or a zinc finger domain.^3^ In this architecture, the chromophore is exposed to nearby residues of the inserted protein. The region between residues 145-148 is where FP barrel slightly bulges towards the solvent to accommodate the chromophore. Crystal structure of GFP (PDB: 1EMA) shows that the side chains of two residues in the bulge flank towards the exterior, disrupting the ordered hydrogen bonding network within the 11 stranded barrel structure. Therefore, this region is permissive to construction of new termini and/or insertion of a new protein without compromising the fluorescence. Circular permutation and insertion of a sensing domain exposes the tyrosine derived phenol moiety, rendering its pK_a_ sensitive to local changes in the hydrogen bond network. Local protein environment is important for the chromophore’s spectral properties since the denatured FPs or isolated chromophores are not fluorescent.^4^ The chromophore exists in an equilibrium with its tyrosine-derived phenol moiety either in anionic or neutral state. After illumination, phenol hydrogen becomes acidic and is transferred via an internal hydrogen bond network to a nearby glutamate residue, a well-characterized process termed ‘Excited State Proton Transfer (ESPT).^5,6^ The uncharged phenol form of the chromophore absorbs shorter wavelength light (395 nm) and is less fluorescent whereas the anionic phenolate form absorbs light at a longer wavelength (475 nm), displaying much more fluorescence.^4,7^ Shifting the phenol-phenolate equilibrium due to a conformational change in the sensing domain is the working principle of most single FP based biosensors.^8^

The bulge region is exclusively preferred for sensing domain insertion in most of the designs. The two residues flanking the bulge region are called ‘gate post residues’, which are Y145 and H148 in GFP.^9^ Variations in gate post residues as well as the linkers connecting the sensing domain and FPs are diverse among reported biosensors.^9^ During a typical design process, a pool of sensors with randomized gate posts and linkers are screened for the highest ligand dependent fluorescence change. Linkers are kept as short as possible to allow for efficient coupling between the chromophore and the sensing domain, but not so short that folding of either partner is compromised. Defining characteristics of successful biosensors have been reviewed in depth based-on reported crystal structures.^9^ In most cases, the mechanism of allosteric modulation of fluorescence is speculated based on crystal ligand bound structures. It is not clear how the presence or absence of the ligand affects the chromophore environment, specifically the hydrogen bonding network and the phenol-phenolate equilibrium. Crystal structure of ligand-bound sensors reveal that the anionic chromophore is typically stabilized by a hydrogen bond donor located in the linker, sensing domain or the FP itself (Figure 1A-1B).^10–13^ The ultimate effect of the conformational change occurring in the sensing domain upon ligand binding is to move a hydrogen bond donor near the chromophore phenol oxygen which can stabilize the anionic charge formed after the ESPT reaction. This hydrogen bond donor presumably moves away from the chromophore in the OFF state; however, only a few crystal structures of both ON and OFF states of the same biosensor have been reported. ^10,14^ Potassium bound ON state structure of GINKO1 represents an alternative scenario; the basic residue E295 is in close proximity of the chromophore phenoxy oxygen, possibly acting as a proton acceptor in ESPT reaction and providing directional hydrogen bonding to the deprotonated chromophore (Figure 1C). This idea was used to create a long Stokes shift calcium indicator; a positively charged lysine residue which originally stabilized the deprotonated phenolate moiety was mutated into a negatively charged glutamate to act as a proton acceptor.^15^ Alternatively, a basic residue may be positioned near the chromophore which stabilizes the neutral OFF state and moves away in the ON state, as exemplified by the nicotine sensor iNicSnFR1 (Figure 1D).^14^

**Figure 1.**
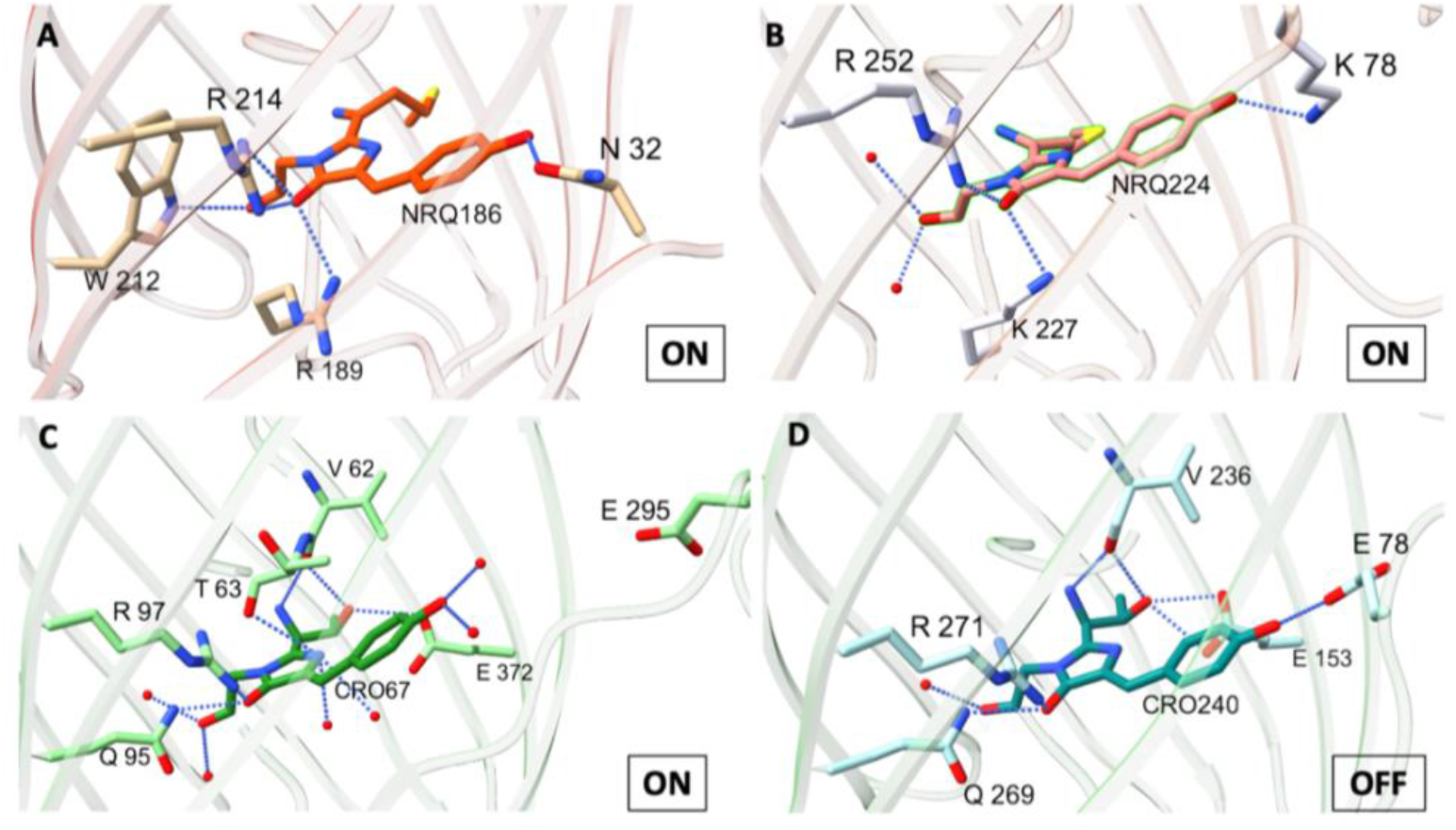
Chromophore environment in ON or OFF states of **A**. K-GECO (PDB:5UKG); **B**. R-GECO1 (PDB: 4I2Y); **C**. GINKO1 (PDB: 7VCM); **D**. iNicSnFR1 (PDB:6EFR). Chromophores are labelled and colored according to the color emitted in their fluorescent states. Chromophore coordinating residues are also displayed and labelled according to the numbering in their PDB structures.

All GEFB designs are based on the shifts in the energy landscape of the polypeptide upon ligand binding. Moreover, most will use locations that allosterically modulate the conformations for insertion of the reporter into the sensing domain, so as not to modify binding site interactions. Efficient sampling of protein conformations and disclosing allosteric regions then becomes a priority in the computational design process.^16,17^

Thus, instead of relying on the single protein conformations for interpreting the functioning of the biosensor, using dynamics to disclose the range of available motions is a more rational approach^18^ and is already being used as a strategy in the computational design of GEFBs. ^19^ With the advent of AlphaFold predicted structures^20^, new designs will inevitably be increasingly proposed, and sampling the functionally relevant conformations of the predictions will also be an issue to consider. For example, a recent report revealed the working mechanism of a single FP based camp biosensor by using metadynamics molecular dynamics (MD) simulations to sample the *apo* form starting from the *holo* crystal structure.^21^ Such dynamical approaches are also effective in predicting the role of environmental conditions such as pH and ionic strength^22,23^ and are essential for interpreting the conformational multiplicity of proteins detected by e.g. SAXS experiments.^24^ On the downside, it is not straightforward to deduce ON and OFF states of the conformations obtained from MD simulations which are based on classical physics, while the proton transfer reaction requires very costly quantum chemistry calculations.^5,25^ It is therefore essential to derive descriptors for conformations obtained from MD simulations to characterize the ON state of a GEFB.

In this study, we examined the dynamics of *holo* and *apo* forms of a maltose biosensor comprised of *E. coli* maltose binding protein (MbP) and circularly permuted green fluorescent protein using classical MD simulations. This experimentally well-characterized biosensor was reported to have a high signal-to-noise ratio, bright enough to visualize maltose transport across *E. coli* periplasmic membrane and extracellular maltose in mammalian cells despite the moderate degree of conformational change.^26^ *E. coli* MbP is a member of bacterial periplasmic binding protein family and consists of two domains linked by a two- or three-stranded hinge. The ligand binding site is found at the domain interface. The hinge bending motion drives open-to-closed conformational transition upon ligand binding.^27^ In this work, we deduce the allosteric effect of maltose binding upon the chromophore environment by analyzing several 1.2 μs long MD simulations of the *apo* and *holo* forms. Our aim is to put forth the descriptors that would best characterize the ON/OFF states of single FP-based nanosensors. We believe the mechanistic insights developed in this work will facilitate the design of novel single FP biosensors in a much shorter time by minimizing the experimental trial-and-error process.

## 2. Methods

### 2.1. Modelling the initial coordinates of *apo* and *holo* sensors

Colabfold was used to obtain an initial structure with completed missing residues using pdb100 template search option and Mmseq2 multiple sequence alignment (MSA).^28^ The sequence of maltose bound sensor named as ‘MbP311cpGFP’ where cpGFP is inserted at position 311 of MbP was used (Figure S1).^26^ Since the chromophore is a non-standard residue that is not recognized by Alphafold, the three-residue-sequence ‘TYG’ was used in the input sequence in place of the chromophore.^29^ The highest ranked model was then aligned with the template PDB: 3OSR structure and the coordinates of the cyclized chromophore were transferred into the model pdb coordinates. For the *holo* state, coordinates of maltose molecule were also added. E390 within the fluorescent protein barrel was modelled neutral as predicted by PROPKA3.^30^ This residue structurally aligns with E222 in intact GFP (PDB:1EMA) which is predicted to be the proton acceptor in ESPT.^31^ Since this sensor produces higher fluorescence in the ligand bound form, a so-called ‘direct response sensor’, we modelled the *holo* state with an anionic chromophore and the *apo* state with a neutral chromophore, representing the ON and OFF states of the sensor, respectively.

### 2.2. MD simulations and postprocessing

VMD and NAMD packages were used to prepare the simulation box and simulate the dynamics of biosensor models in water.^32 33^ The common simulation protocol applied in all runs was as follows: The protein was placed in a rectangular box with at least 10 Å layer of water in each direction from any atom in the system. The simulation box was neutralized with K^+^ and Cl^+^ ions while maintaining an ionic strength of 150 mM. The protein, the water molecules and the ions were modelled using the CHARMM36 all atom force field ^34^. Chromophore atoms were modelled using a combination of CGenFF and CHARMM36 parameters. The topology of the neutral chromophore is used as described in CHARMM36 for residue label “CRO”. The deprotonated chromophore topology was created by using phenoxy group topology in CGenFF. Maltose topology was created with CHARMM-GUI Glycan Reader tool.^35,36^ The RATTLE algorithm was applied and the trajectories were calculated using the Verlet algorithm with a timestep of 2 fs. Long-range electrostatic interactions were calculated by particle mesh Ewald summation method with a cutoff distance of 12 Å. All systems were subjected to energy minimization before running in the *NPT* ensemble at a temperature of 310 K and a pressure of 1 atm for at least 1.2 μs. *Apo* and *holo* runs were performed as four independent replicates starting from the same initial protein coordinates. The coordinates were saved every 2 ps for further analysis. Hydrogen bond occupancies of equilibrated trajectories were calculated using the VMD hydrogen bond and VMD Timeline plugins. The donor-acceptor distance and angle cutoff were adjusted as 3.5 Å and 30º respectively. Hydrogen bond lists obtained by the VMD Timeline plugin were processed with a previously published python script which merges all hydrogen bonds between the atoms of two residues into a single entry.^37,38^ The trajectories were clustered using the CPPTRAJ program with *k*-means algorithm based on C_α_ root mean square deviation (RMSD) values.^39^ Solvent accessible surface areas (SASA) were calculated with a VMD Tcl script with a probe radius of 1.4 Å using Shrake-Rupley alghoritm.^40^

## 3. Results

### 3.1. Structures predicted by Colabfold

Rank1 model obtained from Colabfold has overall high position accuracy per residue, termed Predicted Local Distance Difference Test (pLDDT) score, except for the long loop of cpGFP and the three-residue sequence ‘TYG’ at the position of the chromophore (Figure 2A). The relative position of cpGFP and MbP domains is also well predicted, as evident by low Predicted Aligned Error (PAE) throughout the whole protein (Figure 2B). All five models produced by Colabfold showed substantially higher similarity to the closed *holo* state of MbP (PDB: 1ANF). (Table S1). Using *apo* MbP (PDB: 1OMP) as a custom homology template did not shift the predicted structure towards the *apo* form (data not shown). This observation supports a previous study where the authors showed that Alphafold2 has a bias towards predicting the *holo* state of proteins.^41^ Models also showed varying degrees of similarity to the *holo* sensor structure (PDB:3OSR). Differences in RMSD values to the *holo* sensor is probably caused by the difference in relative position of cpGFP to MbP domain which is also hinted by high PAE in the 3^rd^ and lower ranked models (Figure S2A). Since an initial model close to the *apo* form could not be obtained by Colabfold, both *apo* and *holo* MD runs were started with Rank1 *holo* coordinates, where the *holo* runs additionally contained the ligand maltose.

**Figure 2.**
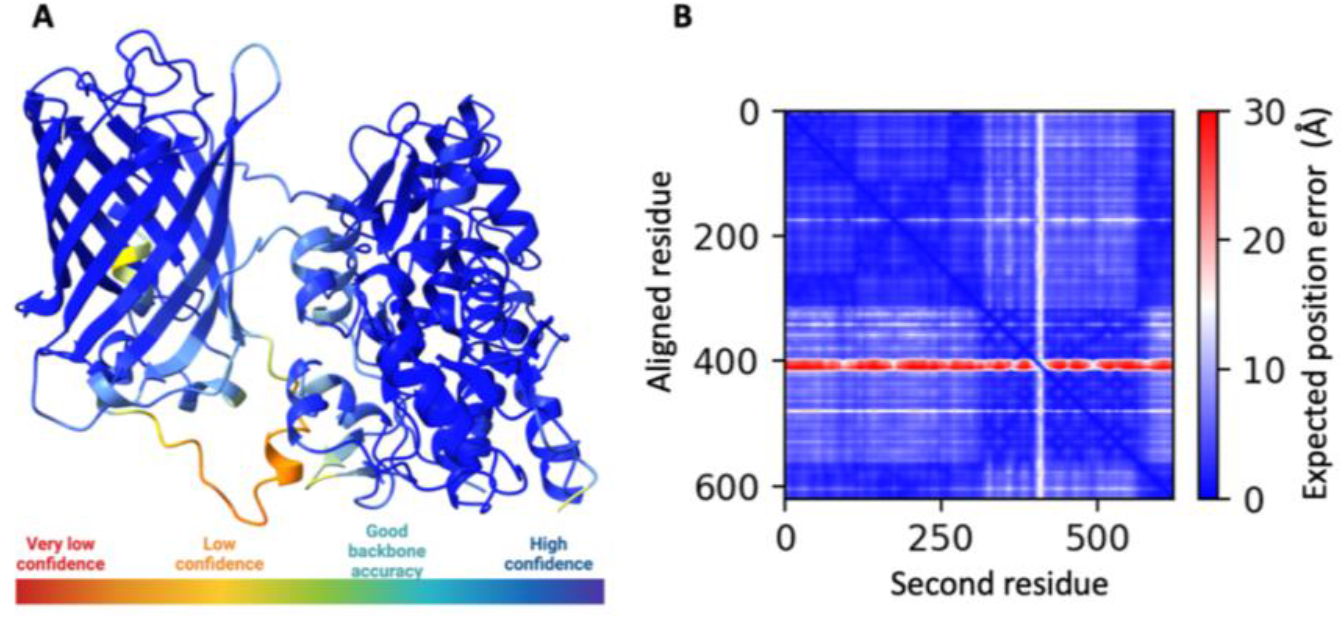
**A**. Rank1 model colored based on per-residue prediction accuracy score. **B**. PAE plot of Rank1 model. Red regions indicate lower accuracy in prediction of relative positions of cpGFP and MbP domains.

### 3.2. MD equilibration and conformational change in *holo-to-apo* transition

Throughout the MD trajectories, the protein preserved its overall conformation in all the *holo* and *apo* runs, with some rotation of FP and MbP domains relative to each other (Figure 3A). In *apo* runs, where maltose was removed from the initial structure, a *holo*-to-*apo* transition was observed. This conformational change is achieved by the outward movement of the residues E45, W63, R66, R67 and W231, all of which are in the N terminal domain (Figure 3A). This finding is in accordance with the previously published observation that the N terminal domain is pulled during the hinge bending movement.^42^ The phenol ring of the chromophore tilted slightly in the *holo*-anionic state, possibly to find a nearby charge stabilizing group (Figure 3B). Coplanarity of the phenol and imidazilinone rings of the chromophore were preserved throughout the trajectory in both the neutral and the anionic states which indicates that there is no destabilization from charged groups within the FP β barrel or the linker region.

**Figure 3.**
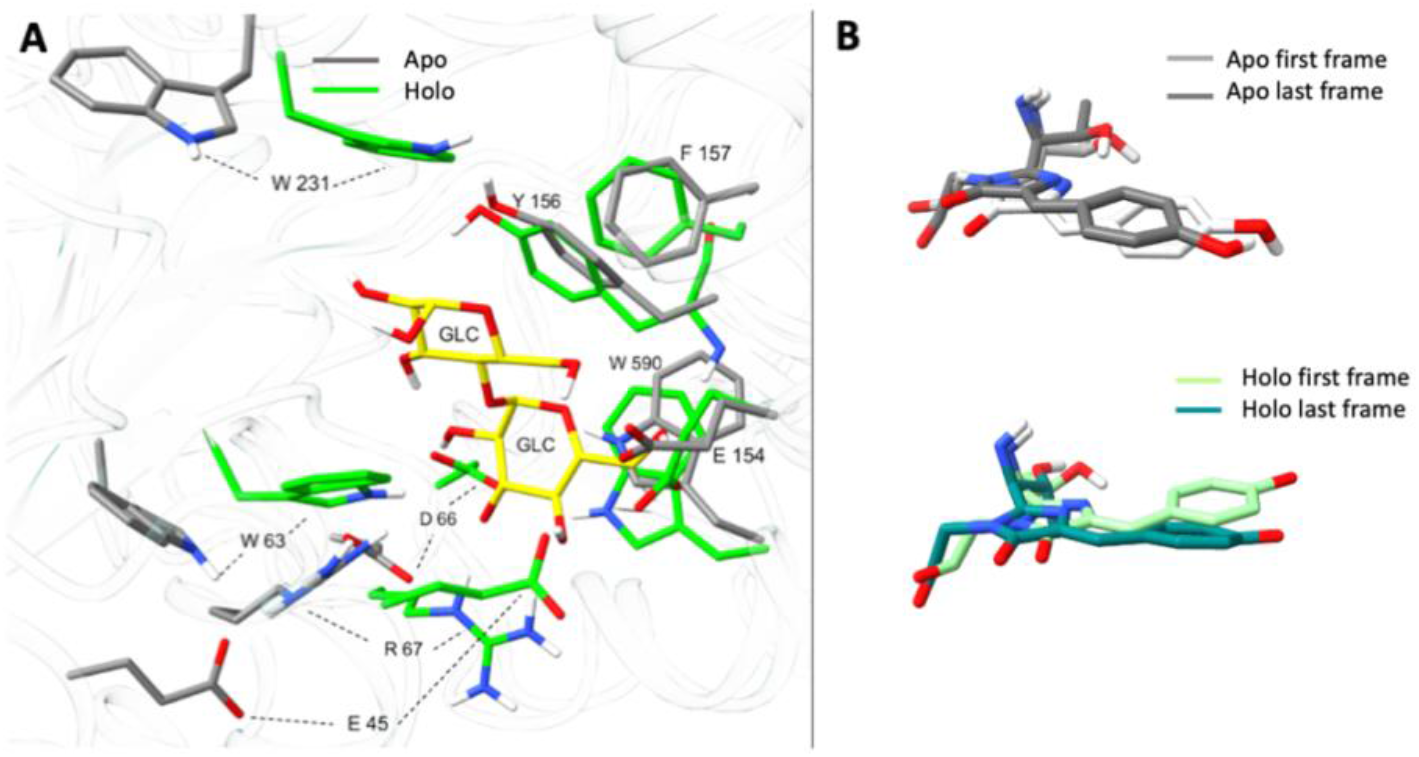
**A**. Maltose binding site in the last frames of *apo* and *holo* runs. Structures are aligned using residues 113-259 and 560-618 covering the MbP domain. **B**. Structural alignment of the chromophore in the first and last frames of *apo*-neutral and *holo*-anionic states. In this and all following figures, green is used for ON (*holo*), and gray for OFF (*apo*) states.

In Figure S3, we display the RMSD of the trajectories and the time evolution of the fraction of the conformational clusters calculated via CPPTRAJ. The regions bounded by the red vertical lines in the RMSD plots indicate the part of the trajectory used in the analyses. To select the snapshots contributing to the equilibrated state in each run, a cluster analysis using CPPTRAJ with *k*-means algorithm was also performed. To obtain a more homogenous population for analysis, we have used the part of each trajectory where one population (pop0) dominates. For all three *holo* runs, pop0 began to dominate the population after 600 ns and remained dominant after that point. For *apo*-1 and *apo*-2, pop0 dominated after 600 ns and 400 ns respectively. In *apo*-3, pop0 dominated between 200 ns-1000 ns, after which time point another cluster emerged (pop1). This time point also coincided with a distinct jump in the RMSD plot. We have extended this run to 1600 ns. At this time point pop1 continued to increase (data not shown); therefore, for this run, we used the section between 200 ns-1000 ns for further analysis. Similarly, for *holo*-4 run, the dominant conformation was sampled between 200 ns-800 ns.

### 3.3. C_α_ root mean squeare fluctuations (RMSF)

RMSF values were calculated using the equilibrated part of each trajectory. Average fluctutation values are higher in the cpGFP domain in both *apo* and *holo* runs (Figure 4A). This trend is also reflected in B-factors of the *holo* sensor crystal structure. RMSF of cpGFP residues increase even further in the *holo* state, where RMSF of MbP residues are comparable in both states. Lower RMSF values for cpGFP domain in the *holo* state may reflect a general property of proteins that was previously revealed, where it was shown that while the binding site residues become more rigid upon ligand binding, flexibility of distant residues tends to increase.^43,44^ In line with this observation, the average B-factor value of maltose-bound crystal structure of MbP is higher than that of the maltose-free structure, except for the binding site residues.^42^ The last frame of the *apo*-1 run has a backbone RMSD of 2.2 Å and 2.3 Å when aligned with *apo* and *holo* MbP crystal structures, respectively. Although the structures obtained at the end of the MD runs do not fully reflect the open unliganded form of MbP, compared with the RMSD values presented in Table S1, we find that it deviates from the *holo* form initially obtained by Colabfold and approaches to an *apo*-like state.

**Figure 4.**
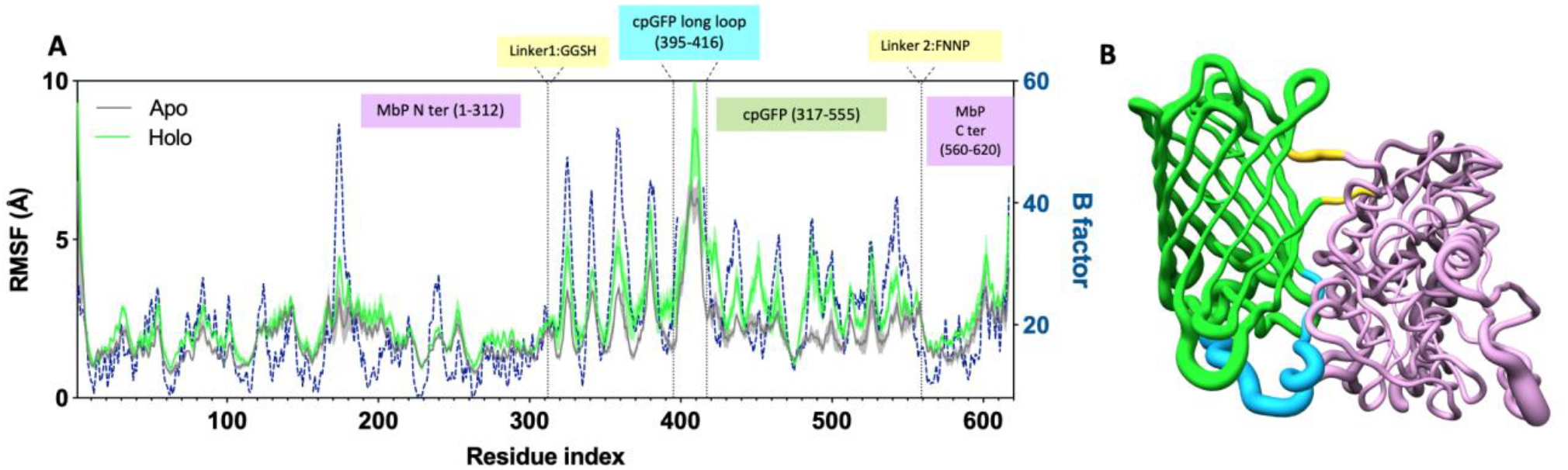
**A**. C_α_ RMSF of the protein calculated over the equilibrated trajectories of *apo* and *holo* runs. Standard error of the mean (SEM), shown as shaded regions, are calculated using the mean values of the replicates. B-factors are taken from *holo* sensor crystal structure (PDB. 3OSR). **B**. Worm style cartoon representation of the *holo* sensor. Thicker regions indicate higher RMSF. Color code indicates different regions of the protein and is the same as the region labels in A.

### 3.4. Hydrogen bond dynamics

To understand the dynamics of allosteric modulation of the chromophore environment by the conformational change in the sensing domain, we have analyzed the hydrogen bond occupancies of the whole protein over the equilibrated part of each trajectory. Hydrogen bond pairs whose occupancies change between *apo* and *holo* states by more than 30 % consistently in all four replicas are listed with the corresponding standard errors. These hydrogen bond pairs are grouped according to their locations; those in the MbP domain, those in the cpGFP domain and the chromophore environment covering both linkers (yellow regions in Figure 4B).

#### MbP domain

In the MbP domain, residues with altered hydrogen bond occupancies are concentrated around the maltose binding site (Figure 5A and Figure 5C). *Holo*-to-*apo* transition increases hydrogen bond occupancies between E45-R67, D66-R67, ALA64-ARG67, K43-K47 and R67-W63, all of which originally interacts with maltose in the *holo* form. Re-positioning of these residues can be seen when the last frames of *apo* and *holo* trajectories are aligned (Figure 3A). Hydrogen bond pairs with higher occupancy in the *holo* state are mostly located on the C terminal domain (Figure 5D). Occupancy of the S264-V262 hydrogen bonded pair which is on the central hinge also drops in the *apo* state, presumably due to the hinge twist motion in *holo*-to-*apo* transition.

**Figure 5.**
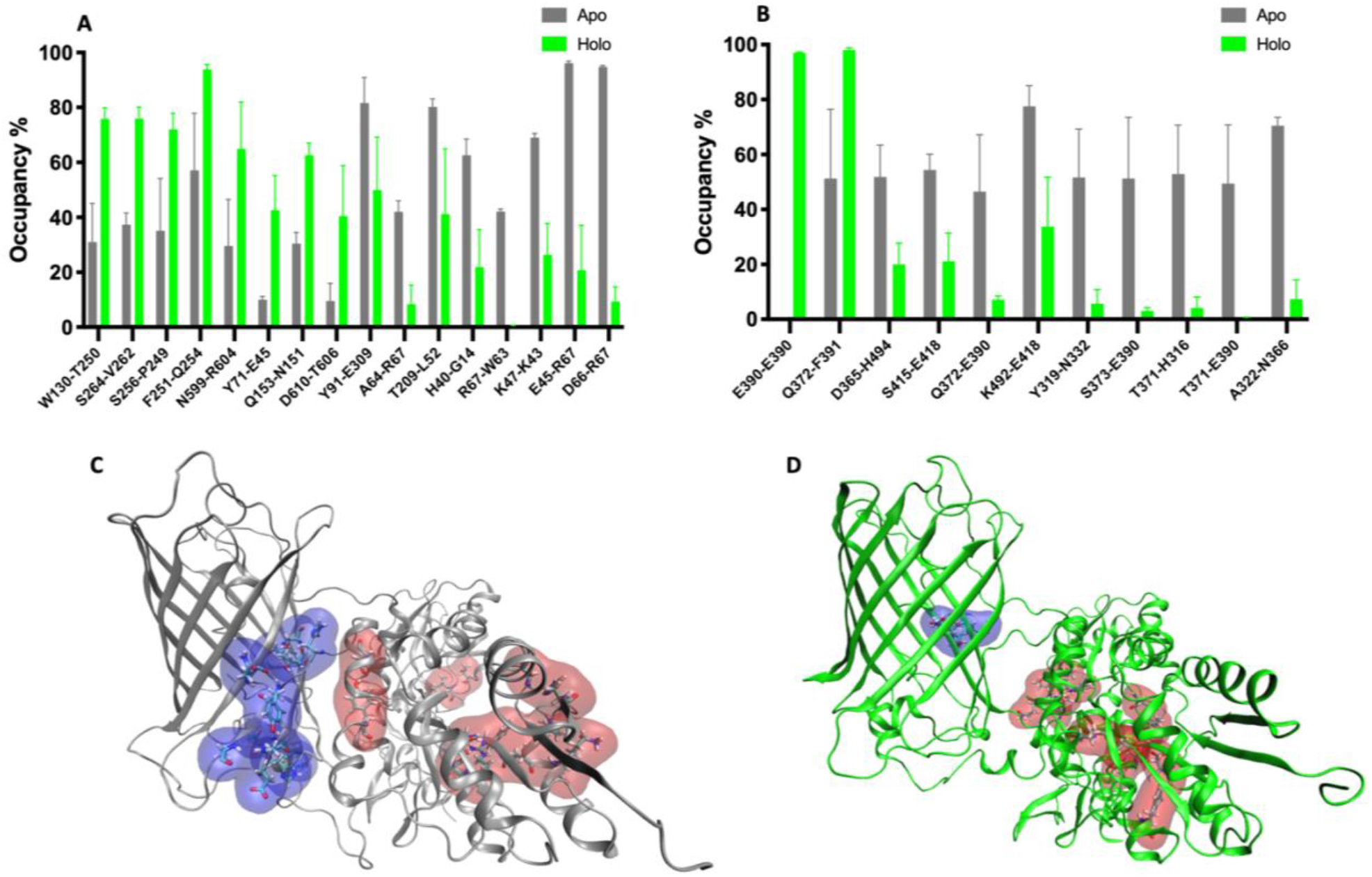
Hydrogen bond occupancies which increase by more than 30 % in *apo* and *holo* state compared to one another. **A**. MBP domain **B**. cpGFP domain. SEM is calculated using the mean values of four replicates for each state. **C**. Location of hydrogen bond pairs with increased occupancies in **C**. *apo* and **D**. *holo* states. Blue and red surfaces display the position of hydrogen bond residue pairs in cpGFP and MbP domains, respectively.

#### cpGFP domain

In the cpGFP domain, the occupancy of majority of hydrogen bonds remains unchanged between the *holo* and *apo* forms. Residues that respond to this conformational change are exclusively located in the two β strands, where the chromophore protrudes through, covering residues 316-323 and 363-376 as well as the loop residues 415-418 and 492-496 (Figures 5B and 5C). These residues constitute the majority of interaction site between cpGFP and MbP. The shifts in the hydrogen bond occupancies are predicted to be the result of the outward bending of the N-terminal (sensing) domain upon ligand removal and re-orientation of cpGFP domain relative to the MbP domain.

#### Linkers and the chromophore environment

Since the pKa of the chromophore is affected by nearby charged residues ^8^, we examined the occupancies of all hydrogen bonds involving the chromophore as either acceptor or donor. Furthermore, nearby residues located in the linker or MbP domain may contribute to either direct or water mediated hydrogen bonds with chromophore phenol oxygen which protrudes into to the sensing domain. We found that the chromophore is hydrogen bonded to several residues from within the β barrel with similar bond occupancies in *apo*-neutral and *holo*-anionic state (Figure 6A). Most notable difference is the increased occupancy of the CRO-T371 bond in *holo*-anionic state. The interatomic distance between the side chain oxygen of T371 and the chromophore phenoxy oxygen is significantly shorter in the *holo* forms (Figure 6B and Figure 6C). T371 structurally corresponds to T203 in intact GFP (PDB: 1EMA); previous MD studies have shown that this residue is hydrogen bonded to the chromophore phenyl oxygen in the ON (anionic) but not in the OFF (neutral) state. ^45, 46^ Other residues which are hydrogen bonded to the imidazolidone ring are V474, Q507 and R509 with similar occupancies. Corresponding residues in intact GFP were also found to be equally involved in hydrogen bonds with the chromophore in both ON and OFF states in the same study.^45^ E390-CRO bond has higher occupancy in the holo-anionic state; in contrast, E222 in intact GFP was found to interact with the chromophore in ON and OFF states with equal geometry.^45^

**Figure 6.**
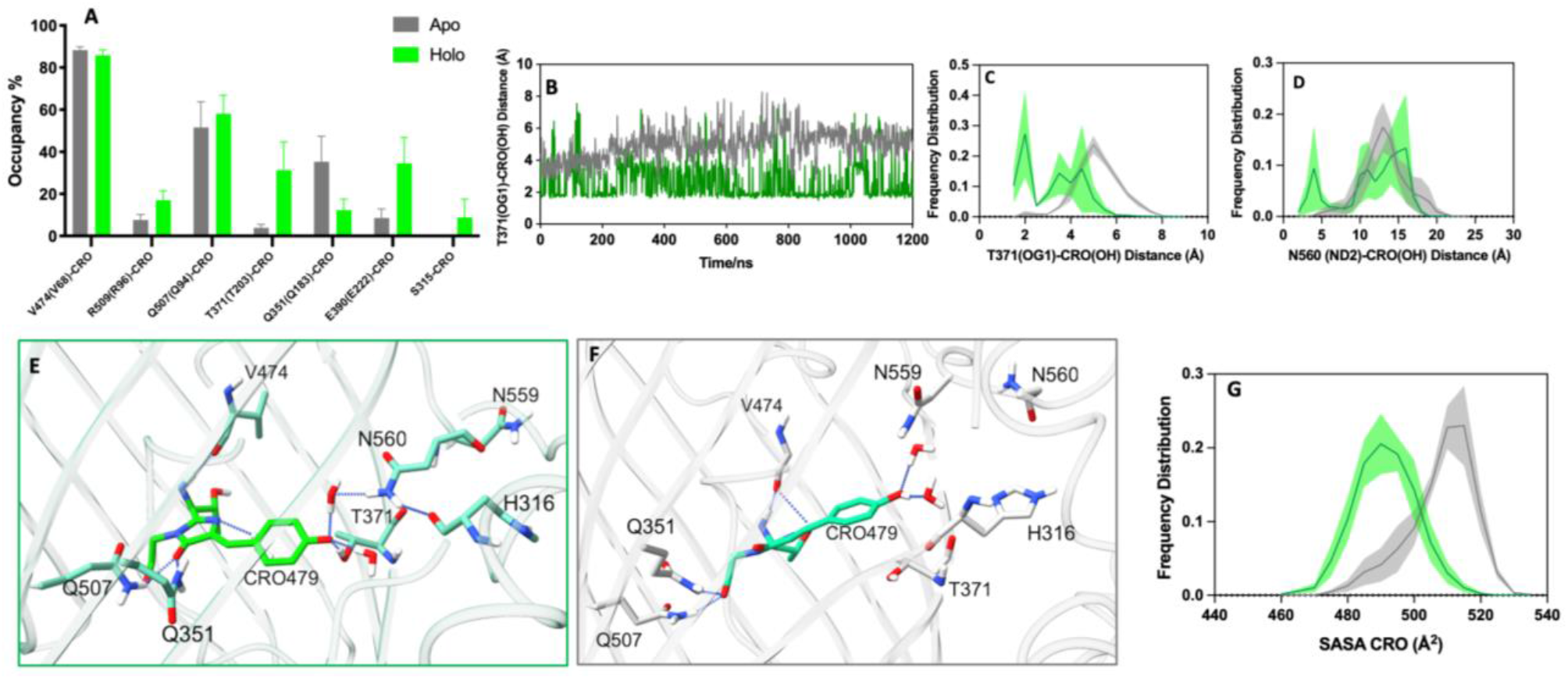
**A**. Occupancies of hydrogen bonds involving the chromophore as either acceptor or donor and within linker residues. Corresponding residue numbers in intact GFP are given in parantheses. **B**. Interatomic distance between side chain oxygen of T371(OG1) and CRO(OH) over *apo*-1 and *holo*-1 trajectories. Frequency distribution of **C**. T371(OG1)-CRO(OH) distance **D**. N560(ND2)-CRO(OH) distance. Chromophore enviroment in the last frame of **E**. *Holo*2 and **F**. *Apo*-2 runs. Hydrogen bonds are visualized with ChimeraX H-bonds utility, with 3. 5 Å and 30 °distance and angle criteria respectively. **G**. Frequency distribution of chromophore SASA in *apo* and *holo* state.

The protonation state of the chromophore phenyl group ultimately determines the fluorescent state of the sensor. Polar residues in the linker region which may affect the phenyl group’s pKa are S315, H316, N317, N559 and N560. We found that the *apo* form backbone oxygen of H316 on linker 1 is hydrogen bonded to the backbone nitrogen of T371 with an average occupancy of 53 % (Figure 5B). In *holo*2 and *holo*3 runs, H316 in the *holo* state prefers to associate with N560 on linker 2 with 52 % and 21 % occupancies, respectively. This interaction positions N560 at a suitable distance from the chromophore to form a water mediated hydrogen bond with the phenoxy oxygen. Frequency distribution of this distance shows that the side chain nitrogen of N560 samples distances less than 5 Å to the chromophore phenoxy oxygen in the *holo* state (Figure 6D). In all the runs, we observed that N559 and N560 readily switch positions, one pointing towards the chromophore and other towards the solvent. In one *holo* run, the side chain of S315 on linker 1 makes a direct hydrogen bond with the chromophore phenoxy oxygen with 35 % occupancy (Figure 6A). Representative hydrogen bond networks around the chromophore in *holo* and *apo* states are displayed in Figure 6E and Figure 6F respectively.

We found that the cumulative effect of altered hydrogen bond occupancies in cpGFP and MbP domains and in the linkers is a network of hydrogen bonds which communicate the conformational change in the ligand binding site to the chromophore environment. In the *holo* state, this network covers the linker region by the hydrogen bond between H316 and N560 which are located on opposite linkers on each side of the bulge. This may help protect the phenolate state of the chromophore in the *holo* state. In the *apo* state, the linker area is visibly more open which increases the solvent accessibility of the chromophore, promoting the low fluorescent state (Figure 5C and Figure 5D). To assess its exposure to the solvent, we measured the SASA of the chromophore in each trajectory. A significant difference between the *apo* and *holo* run SASA values emerges (Figure S4 and Figure 6G). In the *apo* state, the chromophore is more exposed to the solvent with an average SASA of 506 ± 2 Å^2^ and can be more easily protonated by water and lose fluorescence, whereas in the *holo* state the chromophore is more protected with an average SASA of 491±2 Å^2^ and can maintain the anionic charge. The former is a symmetric distribution while the latter is skewed. This is expected for the *apo* form since the initial structure is obtained by removing the maltose from the *holo* prediction, hence leading to longer equilibration times. Thus, the measured 15 Å^2^ difference in the average SASA values is expected to be a lower bound. Our results suggest that the change in the chromophore SASA can be good indicator of biosensor performance. We hypothesize that too small a difference between the *apo* and *holo* states may indicate a low efficiency of the sensor, although this conjecture must be supported by further MD simulations of different biosensors with varying degrees of conformational change.

## 4. Discussion

In this study, we described the molecular mechanism underlying the allosteric modulation of fluorescence of a genetically encoded maltose biosensor. MD runs each totaling ca. 5 μs for the *apo* and *holo* structures reveal a number of aspects of the working mechanism. Firstly, the bright phenolate moiety of the chromophore is stabilized by a direct hydrogen bond with cpGFP residue T371 and partially by a water mediated hydrogen bond with linker residue N560. Our results confirm the role of T203, structural equivalent of T371 in intact GFP, as a hydrogen bond donor to the anionic chromophore. This residue, being located within the FP barrel, is possibly not directly affected by the conformational change in the MbP domain but is rather attracted to the negative charge of the chromophore in the *holo* state. In contrast, N560 is visibly shifted closer to the chromophore due to the repositioning of the linker segment as part of the hinge opening of MbP. In the dark *apo* state, there is no residue close enough to make a direct or water mediated hydrogen bond with the phenol oxygen. Secondly, we found that *apo* and *holo* states are distinguished by the dominance of certain hydrogen bond pairs over the whole protein. Shift in hydrogen bond networks has been shown to contribute to allosteric regulation of many protein systems.^37,47,48^ To our knowledge, this is the first study that sheds light on the working mechanism of a single FP biosensor using hydrogen bond dynamics and chromophore SASA as fingerprints of bright vs. dark states. Our results indicate that hydrogen bonding between the two linkers in the ON state possibly creates a more closed area at the circular permutation site, evidenced by the reduced SASA values for the chromophore, and helps preserve chromophore in the anionic state. A similar argument was made earlier for GCaMP2 which operates on a much larger calcium-dependent conformational change. Calcium-free vs. calcium-bound crystal structures show a greater opening for the *apo* state at the circular permutation site due to the less extensive interaction between the CaM and cpGFP.^49^ Using classical MD simulations, we were able to sample an *apo*-like state of MbP domain starting with the initial coordinates of *holo* state. This approach provides the opportunity to reveal the mechanism of other reported biosensors with periplasmic binding proteins as sensing domains where only the ligand bound structure is available or for sensor designs whereby structures are predicted via Alphafold. Common features of successful biosensors revealed by MD simulations can be utilized in rational design of new sensors or to improve existing sensors with point mutations with much less labor and cost. In future work we will characterize hydrogen bonding patterns and chromophore SASA in other reported sensor structures so as to test the generalizability of these measures as descriptors for GEFB dark/bright states.

## Acknowledgments

This work was financially supported by TUBITAK project no. 121Z329. We thank Zeynep Berksoz and Ebru Cetin for their contributions in hydrogen bond occupancy data calculations. We thank Ali Rana Atilgan for stimulating discussions.

## Data Availability

Available upon request to the authors.

## Figure Legends

**Figure 1.** Chromophore environment in ON or OFF states of **A.** K-GECO (PDB:5UKG) **B.** R-GECO1 (PDB: 4I2Y) **C.** GINKO1 (PDB: 7VCM) **D.** iNicSnFR1 (PDB:6EFR). Chromophores are labelled and colored according to the color emitted in their fluorescent states. Chromophore coordinating residues are also displayed and labelled according to the numbering in their PDB structures.

**Figure 2. A.** Rank1 model colored based on per-residue prediction accuracy score. **B.** PAE plot of Rank1 model. Red regions indicate lower accuracy in prediction of relative positions of cpGFP and MbP domains. **C.** RMSD values obtained when the models are aligned with the given crystal structures.

**Figure 3. A.** Maltose binding site in the last frames of *apo* and *holo* runs. Structures are aligned using C terminal residues 113-259 and 560-618. **B.** Chromophore in the first and last frames of *apo*-neutral and *holo*-anionic states. In this and all following Figures, green is used for ON (*holo*), and gray for OFF (*apo*) states.

**Figure 4. A.** Cα RMSF of the protein calculated over the equilibrated trajectories of *apo* and *holo* runs. Standard Error of Mean (SEM), shown as shaded, are calculated using the mean values of each replicate. B factor values are taken from *holo* sensor crystal structure (PDB. 3OSR). **B.** Worm style cartoon representation of the *holo* sensor. Thicker regions indicate higher RMSF. Color code indicates different regions of the protein and is the same as in A.

**Figure 5.** Hydrogen bond occupancies which increase by more than 30 % in *apo* and *holo* state compared to one another. **A.** MBP domain **B.** cpGFP domain. SEM is calculated using the mean values of three replicates for each state. **C.** Location of hydrogen bond pairs with altered occupancies in **C.** *apo* and **D.** *holo* state. Blue and red surfaces display the position of hydrogen bond residue pairs in cpGFP and MbP domains respectively.

**Figure 6. A.** Occupancies of hydrogen bonds involving the chromophore as either acceptor or donor and within linker residues. Corresponding residues in intact GFP are given in parantheses. **B.** Interatomic distance between side chain oxygen of T371(OG1) and CRO(OH) over *apo*-1 and *holo*-1 trajectories. Frequency distribution of **C.** T371(OG1)-CRO(OH) distance **D.** N560(ND2)-CRO(OH) distance. Chromophore enviroment in the last frame of **E.** *Holo*2 and **F.** *Apo*-2 runs. **G.** Frequency distribution of chromophore SASA in *apo* and *holo* state. Hydrogen bonds are visualized with ChimeraX H-bonds utility, with 3. 5 Å and 30 ° distance and angle criteria, respectively.

