## Supplementary material for "Allosteric Modulation of Fluorescence Revealed by Hydrogen Bond Dynamics in a Genetically Encoded Maltose Biosensor"

```
1   MRGSHHHHHH GMASMTGGQQ MGRDLYDDDD KDRWGSKIEE GKLVIWINGD KGYNGLAEVG
61  KKFEKDTGIK VTVEHPDKLE EKFPQVAATG DGPDIIFWAH DRFGGYAQSG LLAEITPDKA
121 FQDKLYPFTW DAVRYNGKLI AYPIAVEALS LIYNKDLLPN PPKTWEEIPA LDKELKAKGK
181 SALMFNLQEP YFTWPLIAAD GGYAFKYENG KYDIKDVGVD NAGAKAGLTF LVDLIKNKHM
241 NADTDYSIAE AAFNKGETAM TINGPWAWSN IDTSKVNYGV TVLPTFKGQP SKPFVGVLSA
301 GINAASPNKE LAKEFLENYL LTDEGLEAVN KDKPLGAVAL KSYEEELGGS HNVYIMADKQ
361 RNGIKANFKI RHNIEDGGVQ LAYHYQQNTP IGDGPVLLPD NHYLSTQSKL SKDPNEKRDH
421 MVLLEFVTAA GITLGMDELY KGGTGGSMVS KGEELFTGVV PILVELDGDV NGHKFSVSGE
481 GEGDATYGKL TLKFICTTGK LPVPWPTLVT TLTYGVQCFS RYPDHMKQHD FFKSAMPEGY
541 IQERTIFFKD DGNYKTRAEV KFEGDTLVNR IELKGIDFKE DGNILGHKLE YNFNNPAKDP
601 RIAATMENAQ KGEIMPNIPQ MSAFWYAVRT AVINAASGRQ TVDEDLKDAQ TRITK
```

**Figure S1.** Amino acid sequence of 'MbP311cpGFP' reported by Marvin et al. (2011). In our modelling, we have eliminated the N-terminal sequence indicated in gray since it was not resolved in the crystal structure. Chromophore forming three-residue sequence TYG is highlighted in green.

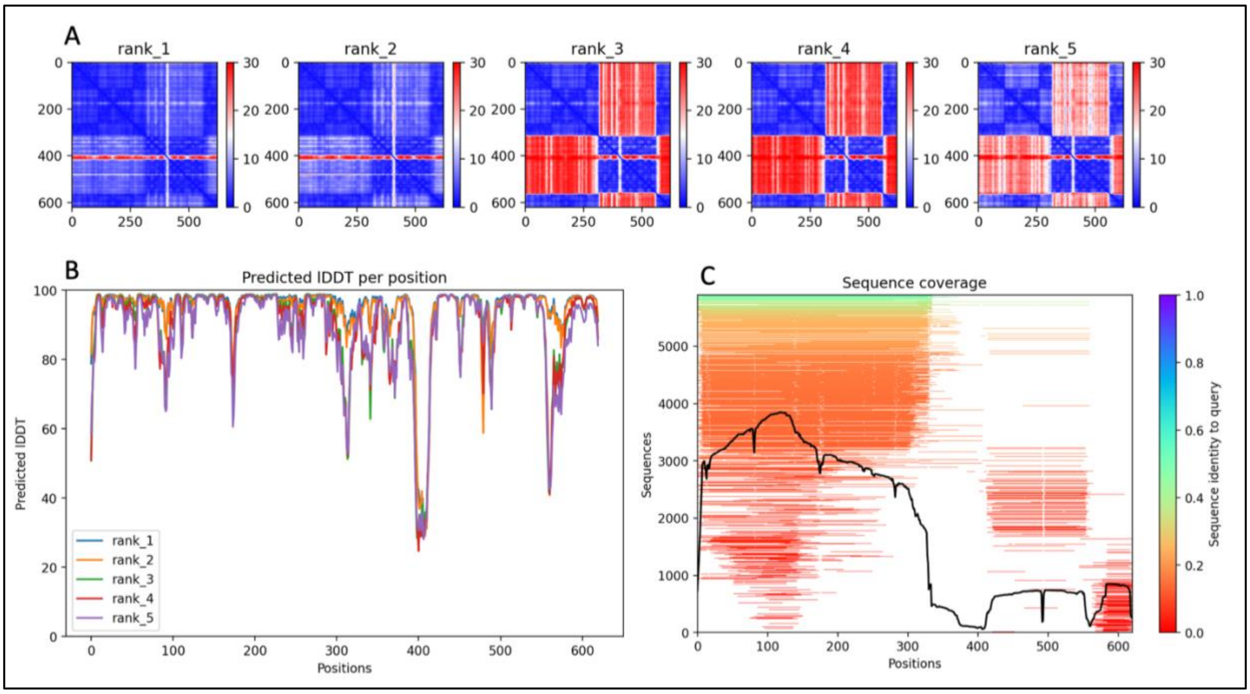


**Figure S2. A.** Predicted Aligned Error (PAE) plots of five models obtained from Colabfold **B.** Predicted Local Distance Difference Test (pLDDT) scores **C.** Sequence coverage obtained from Mmseq2 multiple sequence alignment.

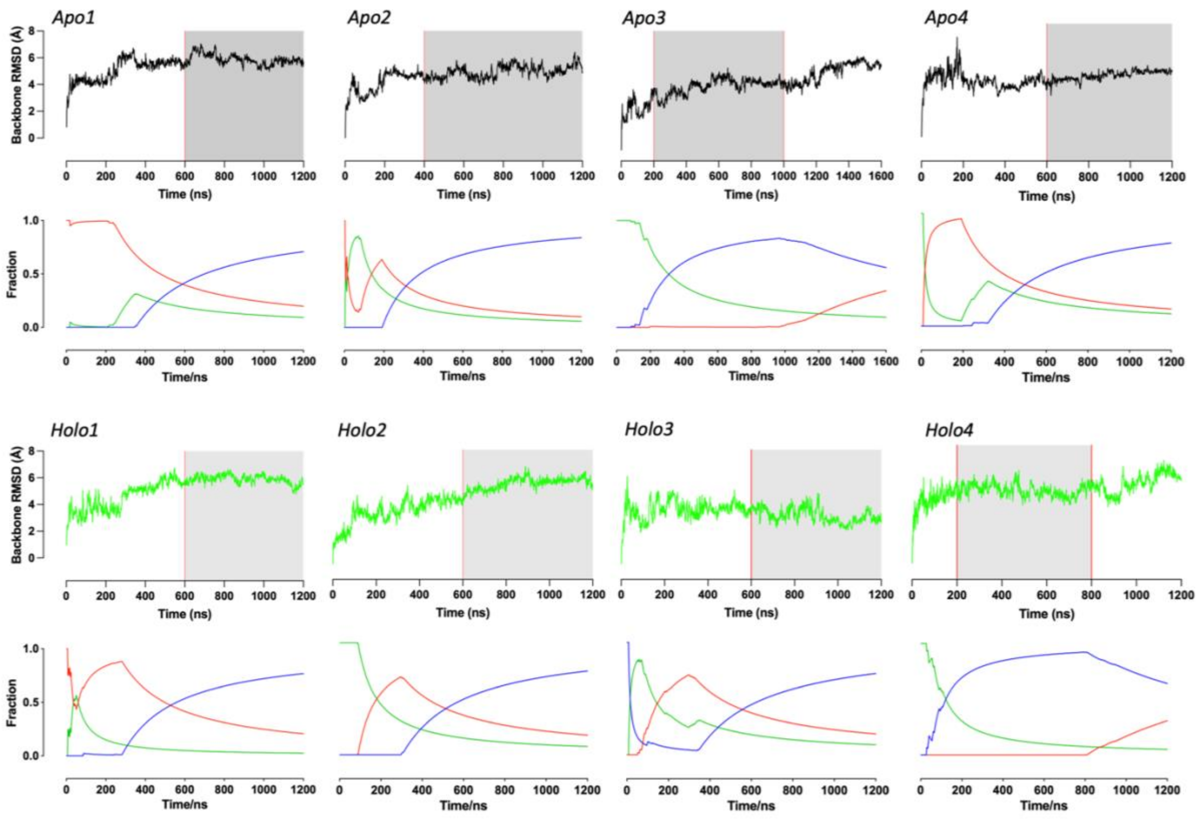


**Figure S3.** Backbone RMSD (upper plots) and Cluster Fractions vs time (lower plots) of *Apo* and *Holo* runs. Part of the trajectories that were used in data analysis is shaded in gray. In cluster vs time plots, **blue, red and green** traces indicate pop(0)-dominant population, pop(1) and pop(2) respectively.

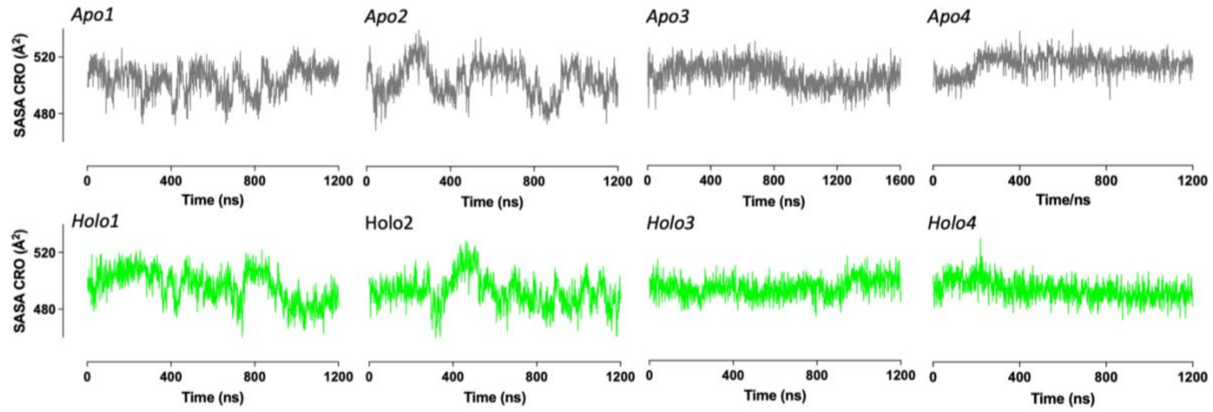


**Figure S4.** SASA of chromophore as a function of simulation time in *apo* and *holo* simulations.

**Table S1.** $C_\alpha$ RMSD values obtained when AF2 models are aligned with the given crystal structures.*

| AF2 model | Backbone RMSD to the template (Å) | | |
|---|---|---|---|
| | Holo sensor (PDB: 3OSR) | Holo MbP (PDB: 1ANF) | Apo MbP (PDB: 1OMP) |
| Rank1 | 0.4 | 0.3 | 3.8 |
| Rank2 | 0.4 | 0.3 | 3.8 |
| Rank3 | 3.3 | 0.3 | 3.9 |
| Rank4 | 1.6 | 0.4 | 3.8 |
| Rank5 | 1.5 | 0.4 | 3.7 |

*Pymol was used to superimpose the structures and calculate the RMSDs. The entire model and the crystal structures listed in the columns were used in alignments.
